# Substrate stiffness regulates collective colony expansion of the social bacterium *Myxococcus xanthus*

**DOI:** 10.1101/2024.05.15.594384

**Authors:** Nuzhat Faiza, Roy Welch, Alison Patteson

## Abstract

Many cellular functions depend on the physical properties of the cell’s environment. Many bacteria have different types of surface appendages to enable adhesion and motion on a variety of surfaces. *Myxococcus xanthus* is a social soil bacterium with two distinctly regulated modes of surface motility, termed the social motility mode driven by type iv pili and the adventurous motility mode based on focal adhesion complexes. How bacteria sense different surfaces and subsequently coordinate their collective motion remains largely unclear. Using polyacrylamide hydrogels of tunable stiffness, we found that wild-type *M. xanthus* spreads faster on stiffer substrates. Here, we show using motility mutants that disrupt adventurous motility suppresses this substrate-stiffness response, suggesting focal-adhesion-based adventurous motility is substrate-stiffness dependent. We also show that modifying surface adhesion by the addition of adhesive ligands, chitosan, increases the amount of *M. xanthus* flairs, a characteristic feature of adventurous motility. Taken together, we hypothesize a central role of *M. xanthus* adventurous motility as a driving mechanism for surface and surface stiffness sensing.

## Introduction

Most bacteria live in surface-dwelling multicellular colonies. Bacteria surface motility plays an important role in various biological and ecological settings, such as microbial infections, the fouling of water systems, and the carbon equilibrium within various ecosystems. Understanding the role of the surface environment in mediating and influencing the motility of microorganisms is therefore central to addressing a wide range of scientific and engineering challenges. The mechanical stiffness of a surface can strongly affect microbe motility, and microbes have evolved various specialized machineries to recognize different surfaces and promote surface exploration. For example, type iv pili are highly conserved appendages that drive propulsion by cyclic extension, attachment, and retraction. The tension on the pili is mediated by surface stiffness, which tunes the cell’s transcriptional response to different surfaces in *Pseudomonas aeruginosa* [1].

In this study, we use *M. xanthus* as a model organism of bacterial collective surface motility. Many microbes have evolved rudimentary forms of multicellularity and collective behaviors, but myxobacteria (δ-proteobacteria) display remarkably sophisticated forms of collective behavior, including swarming, predation, slime trail creation, and multicellular fruiting body development [2-5]. *M. xanthus* possesses two genetically distinct motility systems [6, 7]. The first is called social (S)-motility and is powered by the retraction of type iv pili [8]. The second is adventurous (A)-motility, powered by an inner membrane motor that applies force to the substrate at adhesions [9, 10]. The two motility systems of *M. xanthus* show different selective advantages on different surfaces. When grown on soft agar substrates, *M. xanthus* use S-motility, but on hard agar, *M. xanthus* adopt A-motility modes [11]. These observations suggest that *M. xanthus* adapt and coordinate their motility modes based on physical features of their environment, but the main mechanisms remain unclear.

To determine how bacteria respond to physical changes in their substrate, our group and others have been employing synthetic polyacrylamide (PAA) hydrogels, which have several advantages for testing models of bacteria-substrate interactions compared to commonly used agar surfaces [1, 12-14]. Agar is isolated from marine algae and its isolation process makes it difficult to define and reproduce its chemical properties [15, 16]. It is also difficult to control and tune the physical properties of agar. For example, increasing agar concentration increases substrate stiffness, but also decreases the agar pore size and the ability of nutrients to diffuse into the colony. Further, the viscous properties of agar make it challenging to decipher the mechanics of colony expansion as applied forces will dissipate over time. Polyacrylamide gels on the other hand are linearly elastic with negligible viscous dissipation and linearly deform in response to a wide range of stress. Polyacrylamide gels can be covalently linked with proteins of interest and chemical ligands to the surface to study the integrated effects of specific surface chemistry and substrate mechanics on cell behavior [17].

In this manuscript, we experimentally investigate the effects of substrate stiffness and surface chemistry on the colony expansion in *M. xanthus*. We use polyacrylamide hydrogels of tunable mechanical properties and surface adhesion. We find a range of stiffness over which colony expansion rates increase with increasing substrate stiffness. Using *M. xanthus* mutants deficient in either social or adventurous motility modes, we show this effect is strongly dependent on the adventurous motility mode. Coating surfaces with chitin promotes flare formation and changes the magnitude of expansion but does not change the overall trends with substrate stiffness. Our results underscore the importance of extracellular physical cues on the collective expansion of social bacteria, which has implications for understanding possible mechanosensing strategies used by bacteria.

## Results

### Design and characterization of PAA hydrogels

In the study, we are using polyacrylamide (PAA) gels of varying substrate stiffness and made comparison with commonly used agar gels. Polyacrylamide gels are biologically inert and have well-defined linear elastic properties, making them ideal substrates for studying the effects of extracellular matrix stiffness on cell behavior. We varied the polyacrylamide concentration from 4%-12% and characterized their mechanical properties by performing oscillatory rheology, as we described before [12]. Figure 2 shows representative mechanical data of PAA and agar gels. The data presents the shear storage modulus G’, which characterizes the elastic response of the material, and the loss modulus G’’, which captures viscous dissipation and the fluid-like component of the material. As shown in Fig. 2(a), the rheology of a 2% agar substrate has a significant G’’ value (approximately 6kPa) of the same order of magnitude as its G’ value (approximately 8kPa). In contrast, the PAA gel is linearly elastic with mechanical response dominated by its elastic storage modulus (G’>>G’’).

In some cases, we vary G’ of the PAA gels by increasing the amount of bis-crosslinker while holding acrylamide concentration constant at 8% (Methods) to probe how changes in the pore size of the matrix impact the colony response [12]. The data in Fig. 2(b) demonstrates one of the advantages of PAA gels for understanding how bacteria respond to variations in substrate stiffness, by allowing us to focus on conditions in which the substrate yields a linearly elastic response and diminishes the effect of complex viscoelastic material properties.

### Substrate stiffness increases M. xanthus colony expansion rates

Our experimental protocol consists of observing the growth of *M. xanthus* colonies on the surface of hydrogel substrates with time-lapse microscopy (Methods). Before inoculation, the PAA gels are soaked multiple times in CTTYE nutrient-rich broth, and we deposit a small inoculum of bacteria on the gel surfaces (Fig. 1).

**Fig. 1.**
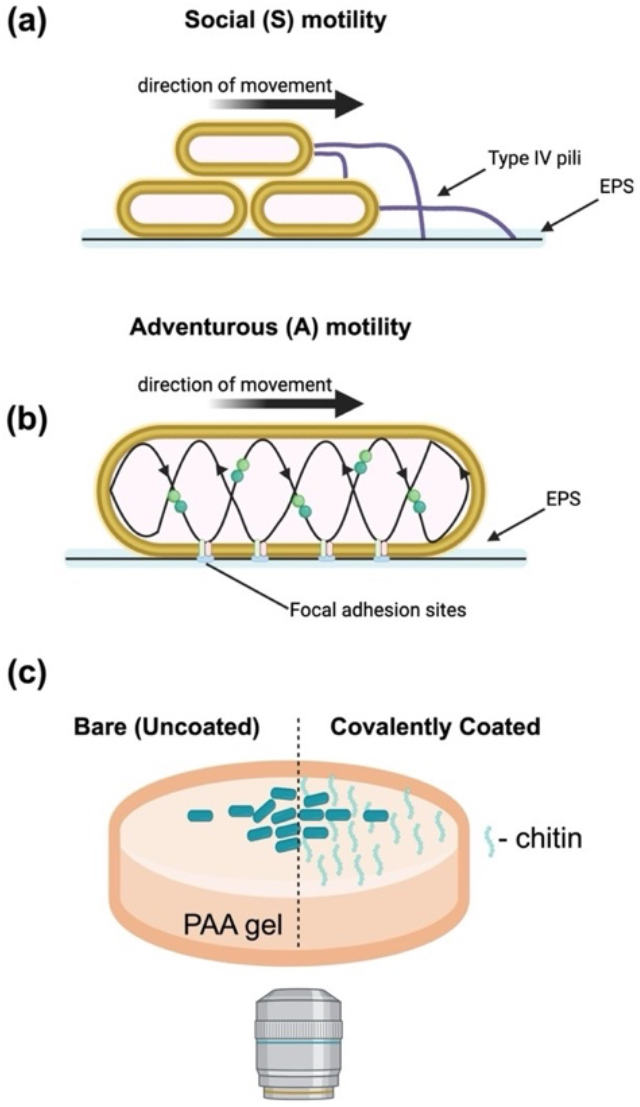
Schematic of the experimental study. *M. xanthus* adhere and move to surfaces using two distinct motility. (a) Social (S) involved attachment and retraction of a surface appendage, type iv pili, that adheres to other cells and cell-secreted EPS trails. (b) Adventurous (A) motility involves secretion of an EPS slime trail and formation of bacterial focal adhesion sites that couple to helically trafficked motor. (c) In this study, *M. xanthus* colonies are cultured and imaged on polyacrylamide (PAA) gels of tunable stiffness and surface conditions. Two experimental surface conditions are used: either (i) inert uncoated gels and (ii) covalently-linked chitin on the surface.

**Fig. 2.**
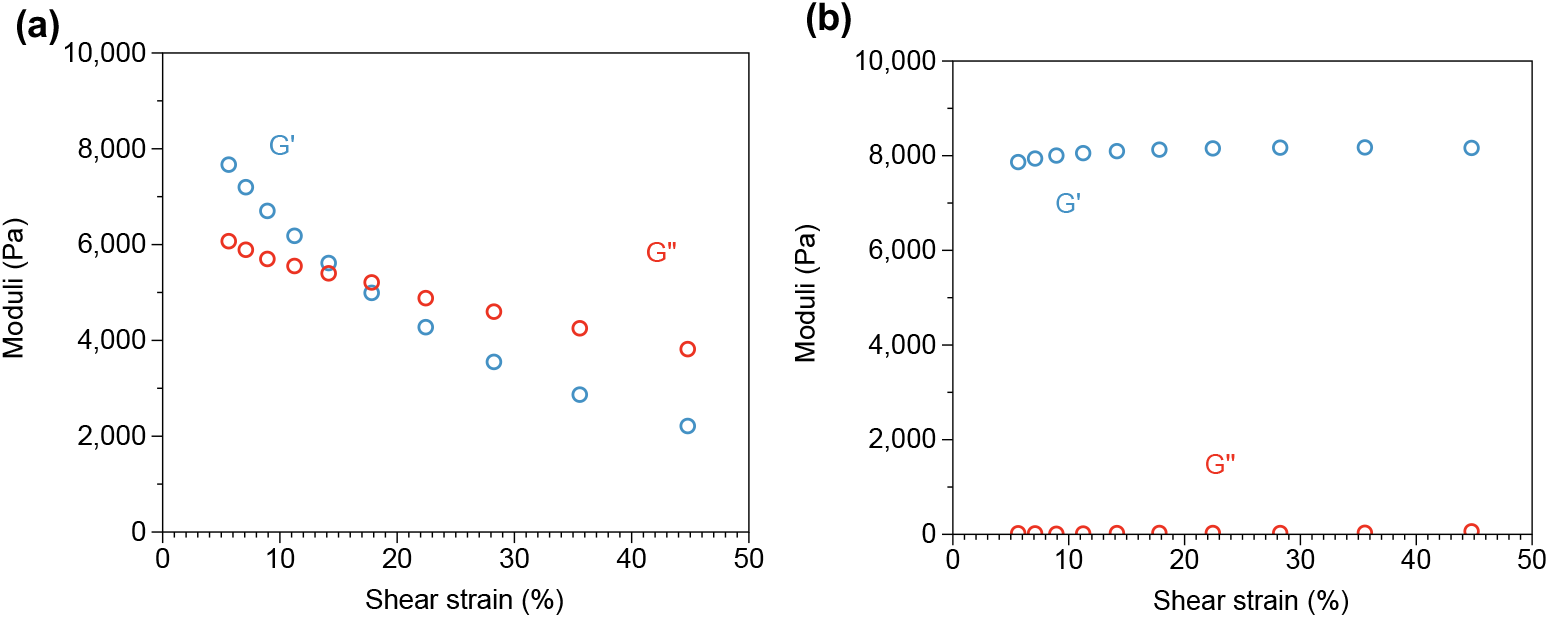
Mechanical characterization of agar and PAA gels. Sample oscillatory shear-strain data from a stress-controlled rheometer of (a) agar and (b) polyacrylamide hydrogel. The plots show the elastic storage modulus, G’, and the viscous loss modulus, G’’, as a function of shear strain amplitude from 5% to 50%. Agar substrates have a significant G’’ value, indicating a strong viscoelastic response to external forces.

Representative images of the *M. xanthus* colony expansion is shown in Fig 3(a). The images show differences in the shape of the colony edge and the colony expansion rates with changes in PAA concentration. SI Table 1 presents the shear modulus for the varying PAA concentrations. In particular, we find prominent protrusions from the colony edge on stiffer PAA gels (5-12 kPa), while the protrusions and edge fluctuations are strongly suppressed on the softer PAA gels (1 kPa). To quantify the colony expansion, we measure the average position of the colony front with respect to its starting location at five-hour intervals and compute the mean colony expansion velocity as a function of PAA concentration. The data show an increase in colony expansion rates with increasing PAA concentration. This enhancement with PAA concentration is somewhat counterintuitive since *M. xanthus* colonies are known to have slower expansion rates on more concentrated (and stiffer) agar substrates. Indeed, this same species the colonies are smaller on more concentrated agar [11, 18] (SI Figure 1). The data in Fig. 3(b), (c) is thus unexpected.

**Fig. 3.**
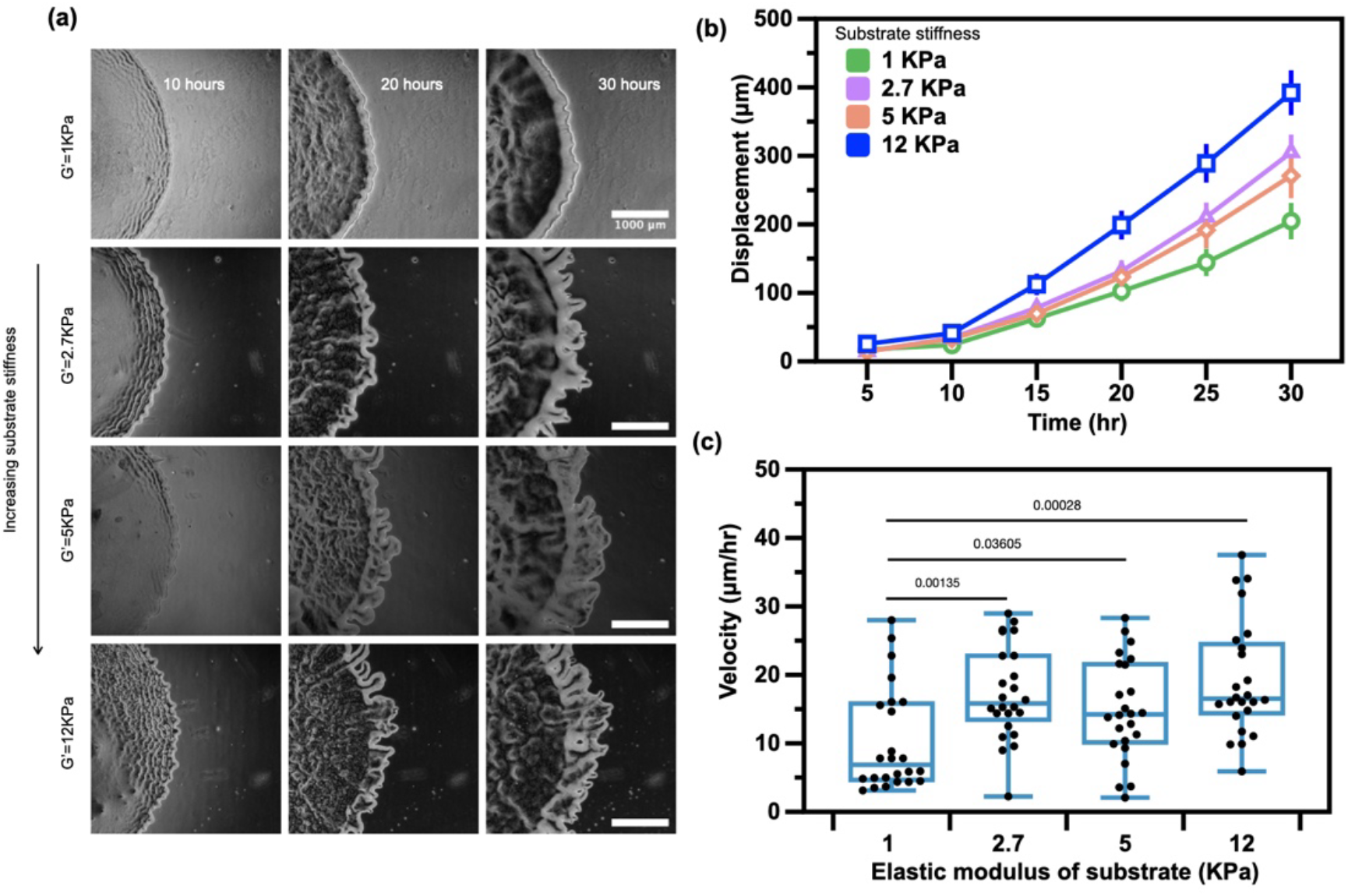
*M. xanthus* wildtype colony expansion on PAA surfaces of varying stiffness. Qualitative and quantitative comparison of colony edge expansion phenotype on PAA substrates of varying stiffness. (a) Representative images of qualitative differences in colony edge expansion phenotypes of *M. xanthus* DK1622 (WT) on substrates of increasing (top to bottom) stiffness at 10, 20, and 30 hrs, after inoculation. (b) Colony edge displacement with time, showing greater colony expansion on stiffer PAA gels. Comparison of colony edge expansion velocity between the (c) 20-30 hrs timepoint.

Similar increases in colony expansion rates with increasing PAA concertation have been previously reported for the bacterium *Serratia marcescens* [12]. It was argued that the increased colony expansion rates originated from a fluid dynamic model involving osmotic swelling of the colony, which has been described for several members of gamma proteobacteria (*E. coli, V. cholera*) and the gram-positive bacteria *B. subtilis* [19-22]. In this model, as cells secrete components of the extracellular polymer matrix, an osmotic pressure gradient is generated between the colony and the underlying substrates, allowing fluid to flow from the substrate into the colony matrix and resulting in increased colony growth. *M. xanthus*, in contrast, moves exclusively on solid surfaces through A- and S-motility modes [23], suggesting a new mechanism behind substrate-dependent colony expansion that we explore further here.

In addition to substrate stiffness, another physical feature of a hydrogel substrate is its pore size (schematic). The substrate pore size can impact the diffusion of nutrients from the substrate into the colony [24] and also has been hypothesized to impact the density of binding sites between the cell and the substrate [25]. Increasing the amount of PAA increases the substrate stiffness but also impacts the pore size of the underlying substrate. To examine whether *M. xanthus* colonies were responding to the changes in substrate stiffness or network pore size, we designed PAA gels of varying stiffness but similar pore size but varying the amount of bis-crosslinker and holding the concentration of PAA the same. Figure 4(b) shows the colony expansion rate as a function of bis-crosslinker. These results are strikingly similar to the effect of increasing substrate stiffness with increasing PAA, namely the colonies showed increased expansion rates on PAA gels of increasing cross-linker, suggest that in the limit of these purely elastic gels, *M. xanthus* is capable of sensing and responding to the stiffness (G’) of the substrate independently of the substrate network pore size.

**Fig. 4.**
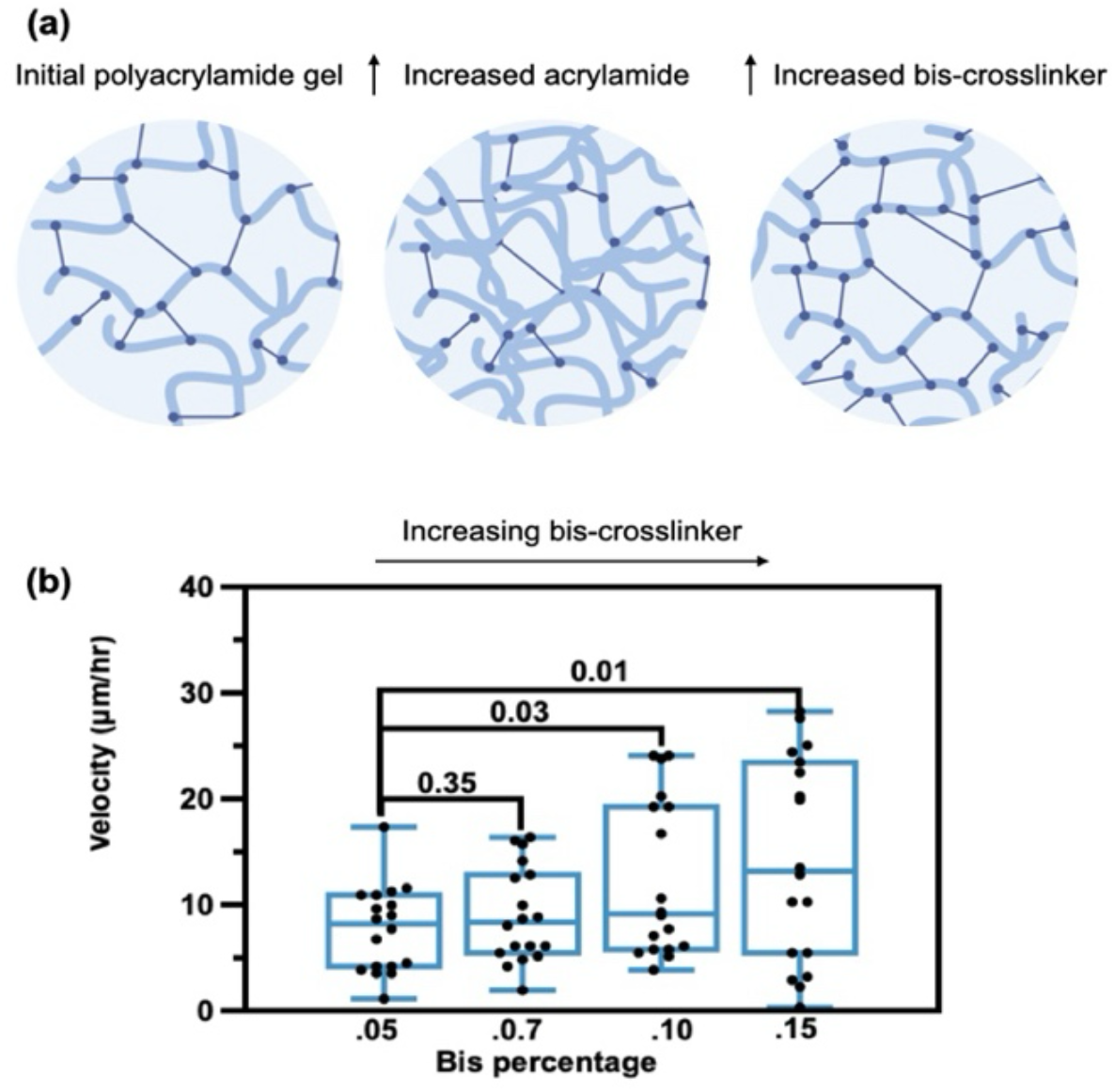
*M. xanthus* wildtype colony expansion on PAA surfaces of increasing bis-crosslinker. (a) Schematic illustration of the effects of increasing acrylamide content versus bis-crosslinker on the hydrogel properties. Both effects increase the hydrogel stiffness, but the impact of bis-crosslinker on the hydrogel mesh size is significantly less than that with increasing acrylamide. (b) Colony expansion rate increases with increasing substrate stiffness via bis-crosslinker percentage. Velocity measured at 10-20 hour time point.

### Effects of A- and S-motility on substrate-dependent response

To connect the wild-type colony expansion to *M. xanthus* motility modes, we next examined the expansion behavior of colonies in species defective in either A motility or S motility (Fig. 5 and Fig.6). Here, we used two historic reference strains, DK1218 (*cglB2*), defective in A motility (A^−^ S^+^), and DK1253 (*tgl-1*), defective in S motility (A^+^ S^−^) [7, 11]. Figure 5 shows representative images of A^−^ S^+^ and A^+^ S^−^ on soft (1 kPa) and stiff (5 kPa) polyacrylamide hydrogels.

**Fig. 5.**
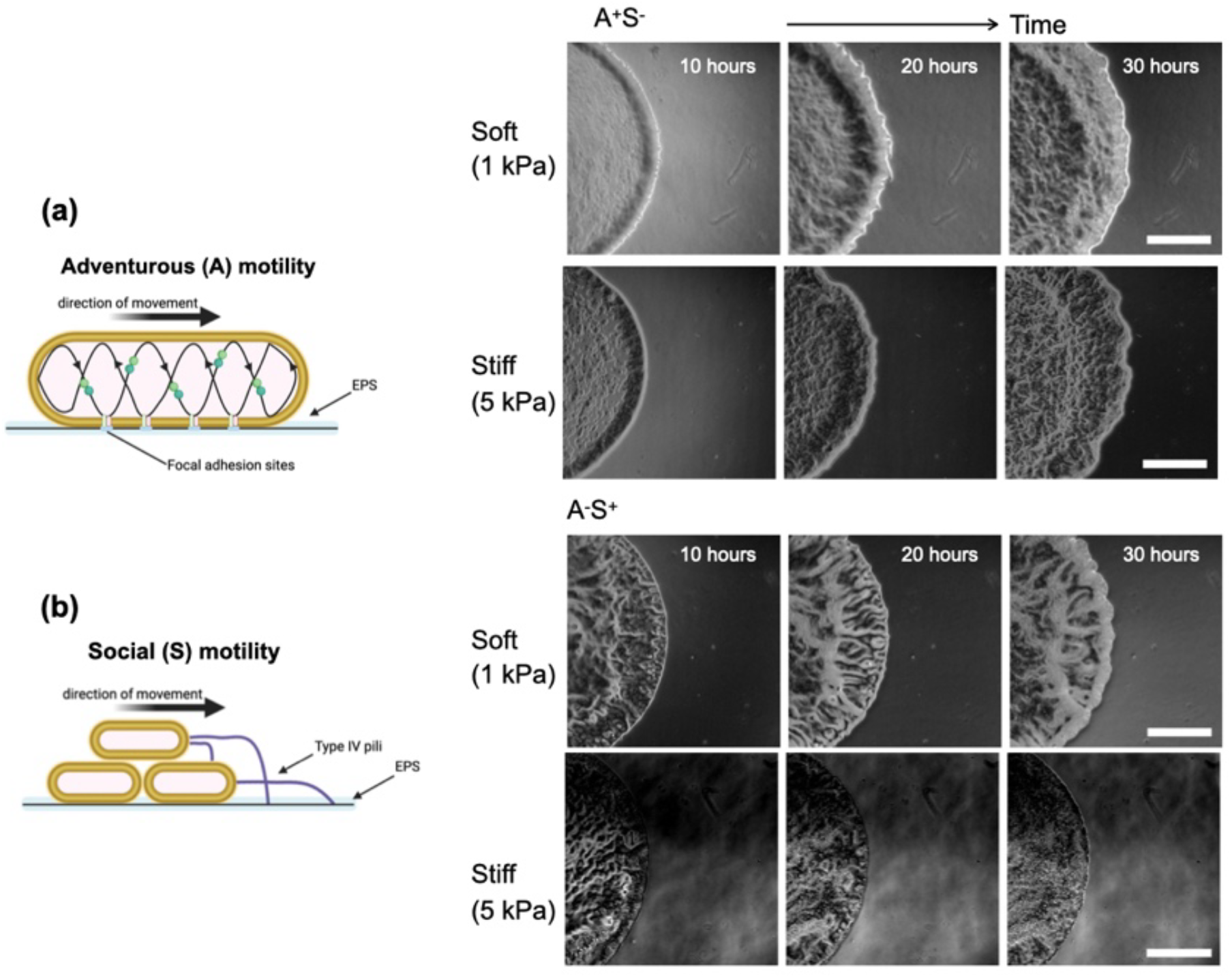
*M. xanthus* motility mutants on soft (1 kPa) and stiff (5 kPa) PAA substrates. Representative images showing the colony edge phenotypes of *M. xanthus* DK1253 (A^+^S^-^) and DK1218 (A^-^S^+^) on (a) soft and (b) stiff PAA substrates after 10, 20, and 30 hrs after inoculation.

Most of the colonies show a rough boundary edge, though the flare-like structures seen in wild-type (Fig. 3(a)) are not observed. The A^−^ S^+^ colony on the 5 kPa gel differs in terms of its colony edge shape, exhibiting a very smooth edge, consistent with the social mode of motility.

The colony edge displacement and velocity for wild-type, A+S-, and A-S+ are shown in Fig.6. The data show that the expansion of A+S-colonies significantly varies with substrate stiffness, increasing mean colonies speeds from 25.98 μm/hr on 1 kPa gels to 68.58 μm/hr on 5 kPa gels. The A-S+ colonies on the other hand show no mean difference in colony expansion rates at a velocity of approximately 6 μm/hr on both surfaces. Interestingly, on the soft gels (1 kPa), we find that the colony expansion rate of the A+S-is similar to wild-type, while on the stiffer gel (5 kPa), we find that the A+S-is markedly faster than wild-type species. Taken together, the data in Fig. 6(c) indicates that the adventurous motility mode of *M. xanthus* is sensitive to changes in substrate stiffness and is involved in increasing colony expansion rates on stiffer substrates.

**Fig. 6.**
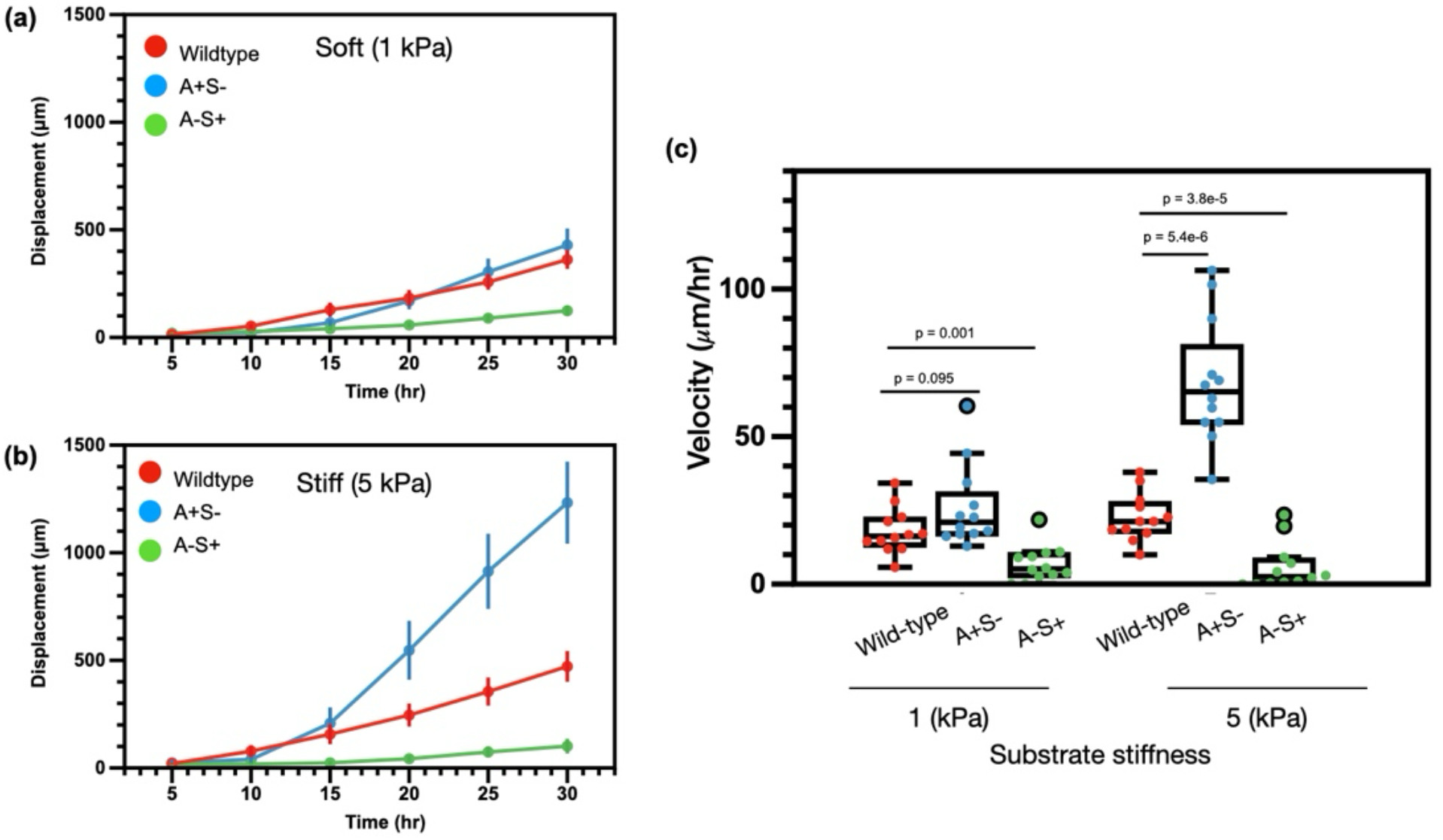
*M. xanthus* wildtype and mutants’ colony expansion on soft and stiff PAA substrates. *M. xanthus* DK1622(WT), DK1253(A^+^S^-^), and DK1218(A^-^S^+^) colony edge displacement with time on (a) soft (1 kPa) and (b) stiff (5 kPa) PAA substrates alongside (c) velocity comparison on both.

### Substrate surface chemistry influences M. xanthus colony expansion

Our experimental data thus far show *M. xanthus* colonies expand at different rates on surfaces of different elastic moduli. Thus far, the colonies have been cultured on bare polyacrylamide hydrogels that are generally considered inert to biological activity and chemical binding [17]. To further investigate this interaction, we prepared PAA gels covalently linked with surface proteins to these otherwise nonadhesive substrates (Methods). In particular, we coated the surfaces with chitin, a natural N-Acetylglucosamine (GlcNAc) polymer, which *M. xanthus* putatively binds to. The specificity of this interaction was highlighted in studies showing the direct binding of pilin protein to chitin required for pilus retraction [26]. We also chose chitin in this study as possible starting point for a ligand-like candidate as *M. xanthus* has putative chitin-binding proteins [27].

Figure 7(a), (b) shows representative images of the wild-type, A+S-, and A-S+ colonies on the chitin-coated surfaces. On the chitin-coated surfaces, prominent flairs associated with *M. xanthus* A-motility can be observed in wild-type and A+S-colonies on the stiff substrates, suggesting that flair formation may be substrate-stiffness dependent. The colony expansion velocities are shown in Fig. 7(c). We observe similar trends with increasing substrate stiffness compared to the uncoated gels, namely, wild-type and A+S-colonies expand faster on stiffer substrates than softer ones, while A-S+ colonies have approximately the same average colony speed on both surfaces.

**Fig. 7.**
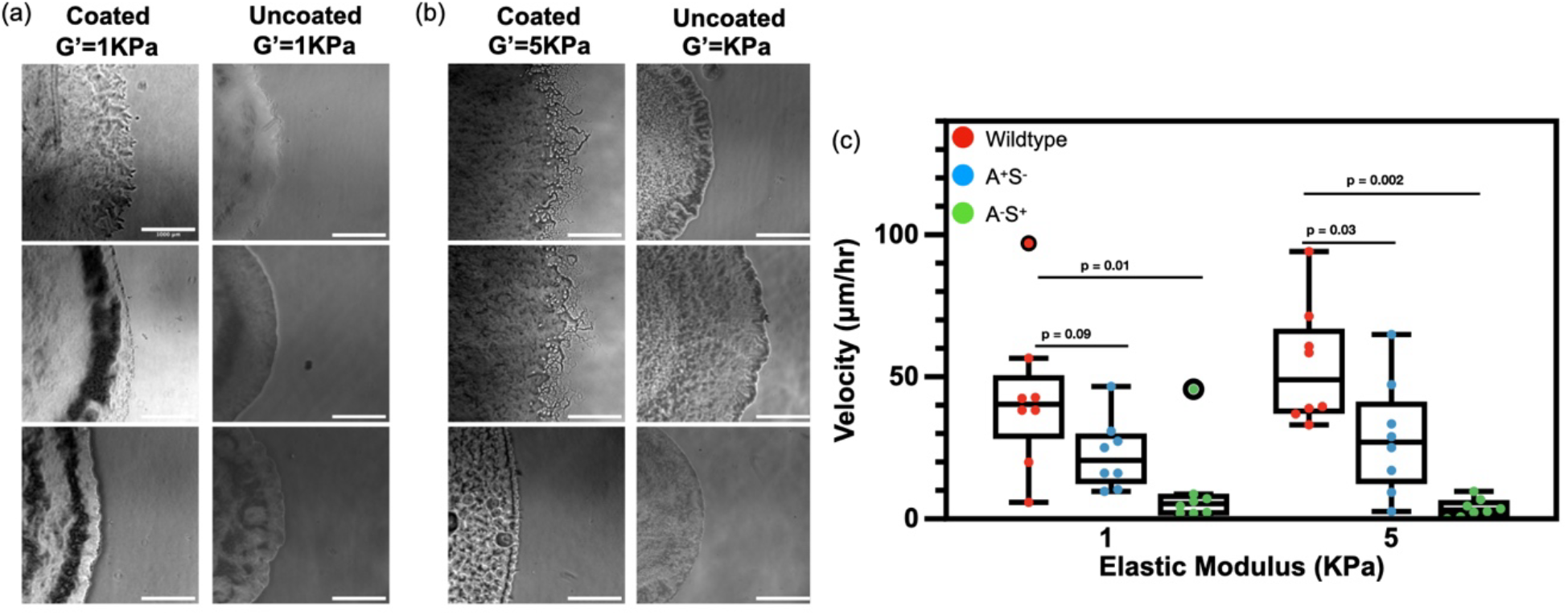
Effects of adhesive protein chitin surface presentation on colony expansion. Representative images of colony edge expansion phenotypes of *M. xanthus* DK1622 (WT), DK1253(A^+^S^-^), and DK1218(A^-^S^+^) on (a) soft (left) and, (b) stiff (right) PAA gels substrates coated with chitin (30 hrs past inoculation). (c) Velocity comparison of colony edge expansion of DK1622 (WT), DK1253(A^+^S^-^), and DK1218(A^-^S^+^) on the coated PAA gels.

One difference between the chitin-coated and bare gels is the relative colony expansion rates between wild-type and A+S-cells. Namely, the A+S-colonies are faster compared to wild-type cells on chitin-coated surfaces, whereas they were slower compared to wild-type cells on the bare gels. Taken together, Fig. 7 shows substrate stiffness impacts wild-type *M. xanthus* and A+S-under conditions where the substrate is chemically modified to present an adhesive protein (chitin) at its surface.

## Discussions and Conclusion

Cell-matrix interactions are a central component of collective bacterial colony expansion. Bacterial colonies change their migration behavior on surfaces with different physical properties, but how they coordinate this process collectively remains largely unclear. Here, we examined the migration of the social bacterium *M. xanthus* on synthetic polyacrylamide hydrogels with tunable stiffness and surface chemistry. We found that *M. xanthus* colony expansion increases with substrate stiffness for both uncoated and chitosan-coated polyacrylamide gels. Further, using *M. xanthus* motility mutants, we found that surface-adhesion-based adventurous motility increased with substrate stiffness, whereas pili-based social motility was less responsive. Our results suggest that the surface-adhesion-based adventurous motility mode of *M. xanthus* enables colonies to modify their expansion on different surfaces and is sensitive to changes in substrate stiffness.

Our study lends new insight into how *M. xanthus* responds to changes in substrate stiffness. Our results indicate that increasing substrate stiffness increases expansion rates of wild-type *M. xanthus* in the limit of purely elastic substrates, unlike conventional agar substrates. These results are consistent with our recent work on *S. marcescens*, which showed colonies respond differently to modifications in substrate stiffness on linearly elastic PAA gels compared to agar substrates [12]. Interestingly, our results are consistent with prior work on *M. xanthus* showing larger colony sizes on stiff agar for S-motility mutants but differ compared to A-motility mutants that were smaller on stiff agar. Our results here underscore the role of substrate stiffness in mediating *M. xanthus* expansion rates and suggest that properties such viscoelasticity and perhaps pore size on mediate colony response on agar.

The strong effect of substrate stiffness on A-motility over S-motility may be surprising given recent work identifying the type iv pili as a bacterial mechanosensing element. For example, in *Pseudomonas aeruginosa*, type iv pili drives surface-specific twitching motility and regulates surface-induced gene expression [25]. Type iv pili allow *P. aeruginosa* to transcriptionally tunes their virulence to surface rigidity through a stiffness sensing mechanism in which retraction of the pili deforms substrate with a force that depends on substrate stiffness [1]. Further, prior work on *M. xanthus* showed pili-based S-motility exert significantly larger local forces on surfaces at 50-100 pN compared to A-motility modes at approximately 1-5 pN [28], which also point to a significant role of type iv pili in generating propulsive forces. Our results here highlight that in addition to type iv pili the focal adhesions formed by *M. xanthus* gliding motility is also important in modifying how *M. xanthus* colonies respond to changes in substrate stiffness. This finding is consistent with prior studies that show the gliding motor is strongly elastically coupled to the substrate [29]. We note that while we find that *M. xanthus* mutants defective in A-motility are less sensitive to changes in substrate stiffness, our results do not exclude a pili-based role in surface sensing. *M. xanthus* expansion engages coordination between both A- and S-motility modes, and A-motility mode involves the deposition of EPS trails that are important for pili-based *M. xanthus* motion [6]. Further, type iv pili mediates cell-cell adhesion, which in the absence of EPS, may be its primary function.

The importance of covalently linking adhesive proteins to the surface of the gels can be found in analogy to the adhesive dynamics of metazoan cells. In metazoans, cell surface attachment proceeds by interaction between specific cell-surface receptors and a chemical ligand. These receptor-ligand interactions can trigger biochemical signals that lead to the formation of focal adhesion sites, which through repetitive assembly and disassembly allow cell migration along a surface. Cells show different behaviors on surfaces of with distinct ligands, because they trigger different biochemical signals. To predict cell migration speeds on surfaces of different stiffness, Chan and Odde pioneered a motor-clutch model of cell migration [30]. In this description, migration is predicted by a model where the rate of adhesion assembly and rate of adhesion disassembly control the cell interaction with the surface and are functions of the tension on the adhesion, which depend on the stiffness of the substrate and the cell pulling forces on that adhesion. *M. xanthus* have been hypothesized to have integrin-like adhesions [31] as part of the A motility gliding machinery, in part based on identification of a von Willebrand A domain-containing outer-membrane protein CglB that couples the gliding transducer at the cell-substrate adhesin sites. Further, a recent model has proposed a mechanism for a substrate-stiffness-dependent migration of an ideal bacterium using type iv pili [32]. In this case, the substrate-stiffness response emerges from the assumption that the binding and unbinding rates of the type iv pili with the substrate is a function of the force on the adhesion, similar to adhesive dynamics in metazoan cells.

Given the generality of receptor-ligand type interactions in surface migration, here we make no assumption a prior on their involvement in *M. xanthus* migration of our gels and simply measure the difference in colony expansion rates on bare and chitin-coated surfaces. Our results show that the presence of chitin on the gel surfaces does modify cell expansion speeds, particularly for the A+S-mutants. These results highlight the importance of surface chemistry in modifying colony expansion and suggest the presence of specific adhesive proteins modify the response of specific motility modes (A-motility). This type of approach combined with experimental measures of the motor-substrate coupling, such as with optical traps demonstrated by Balagam et al [29], will help test and extend models of substrate-stiffness dependent colony expansion. We further note that *M. xanthus* is depositing its own EPS slime, modifying the surface chemistry itself, which presumably is allowing the cells to move on the uncoated bare PAA gels. An interesting next step in future studies would be to isolate specific EPS proteins and covalently-link them to the gel surfaces to identify the adhesive proteins that modify migration.

Taken together, our results indicate an important role of substrate stiffness in mediating the collective expansion of a social bacterium. Our data provides new evidence that the *M. xanthus* gliding machinery modifies colony response on different surfaces and may serve as a potential mechanosensing apparatus. The understanding gained here highlights the significance of mechanical interactions between bacteria and their surroundings as pivotal factors in colony expansion. A natural next step would be to study the influence of substrate stiffness on colony force generation and transcriptional changes.

## Methods

### Cell culture

There were 3 different *Myxococcus xanthus* strains used in this study: *M. xanthus* DK1622 (wild type), DK1253(A+S-), and DK1218(A-S+). To culture, *M. xanthus*, cells were inoculated and grown in CTTYE medium with shaking at 32°C overnight. Cell suspensions were then diluted to an absorption of 1.0 at OD600 in cell medium, and a 5 μL of inoculum was spotted on growth substrates. Cultures were maintained at 32°C for *M. xanthus* for up to 17 hours.

### Gel preparation

Polyacrylamide gels were prepared as described previously [12] of varying stiffness with 4% (G’ = 1000 Pa), 6% (G’ =2700 Pa), 8% (G’ =5000 Pa), and 12% (G’ =12000 Pa) along with 0.15% Bis-acrylamide. Polymerization was initiated by the addition of 1.6 μL electrophoresis grade tetramethylethylenediamine (TEMED) followed by 4.8 μL of 2% ammonium per-sulfate (APS) per 600 μL of final gel solution. For each gel, a total of 200 μL of the solution was then pipetted between two circular glass coverslips, one treated with glutaraldehyde (18 mm in diameter) (bottom) and the other SurfaSil-treated (22 mm in diameter) (bottom). The gels were then allowed to polymerize for 20 minutes. Once the gels were polymerized, the gel and cover slip arrangements were flipped and the SurfaSil treated coverslip was removed from the gels. The final dimensions of the hydrogel formed a disc, approximately 18 mm in diameter and 0.8 mm in height. To prepare PAA substrates for inoculation, protocol previously described by Tuson *et al* [13] was followed. The PAA gels were washed three times (quick wash, 10-minute wash and overnight wash) with TPM buffer. The washes were then repeated with CTTYE medium. Before inoculation, the substrates were removed from the growth medium and dried for 20 minutes with UV sterilization.

### Substrate coating

Two different types of coating were used for the gels, using protocols adopted from prior work [28] The gels were coated in two steps: (1) first, gels were coated with Sulfo-SANPAH (sulfosuccinimidyl 6-(4’-azido-2’nitrophenylamino) hexanoate); Sulfo-SANPAH was mixed with DI water by vortexing in a 15mL centrifuge tube. Then, 1 mL of the mixture was pipetted on top of each gel and Sulfo-SANPAH was allowed to activate by placing the gels under a UV lamp for 20 minutes. In the second step, gels were coated with either chitosan at 66.67μg/mL or DNA (Deoxyribonucleic acid sodium salt from salmon testes) at 100 μg/mL in PBS. Chitosan for coating was prepared first by dissolving 10 mg chitosan in 3 ml of 0.2 M acetic acid, then diluted to a final ratio of 1:50 with DI-water. Coatings were placed on gels and incubated at room temperature for one hour. The gels were then washed with CTTYE following the same three step washes.

### Rheological characterization

Rheology measurements were performed on a Malvern Panalytical Kinexus Ultra + rheometer equipped with a 20 mm diameter plate. The elastic gel solutions were polymerized at room temperature between the rheometer plates at a gap height of 1 mm (30 minutes). The shear modulus was then measured as a function of shear strain from 5% to 50% at a frequency of 1 radian/second.

### Imaging

Time-lapse imaging was performed with a Nikon Ti-E inverted microscope equipped with a 4x objective and the Leica DMi8 inverted microscope equipped with a 4x objective. The cultures were maintained at 32°C for *M. xanthus* using a Tokai-Hit stage top incubator. Phase contrast images were taken every 10 minutes for 30 hours using a motorized stage to capture growth at two positions along the edge of each biofilm. For each colony, two videos were taken at different point locations of the colony edge.

### Analysis and Statistical Methods

The colony edge displacement was measured manually using Fiji-ImageJ. For each colony, two videos were analyzed from two different locations of the colony edge, and at least two technical replicates of each condition were completed per experiment (yielding 4+ measurements per condition per experiment). Independent experiments were repeated at least twice on two different days.

## Acknowledgments

We thank Salim Islam for their insightful discussions. This work was supported by NSF MCB 2026747 award to A.E.P.

