## SupplementalData for "Substrate stiffness regulates collective colony expansion of the social bacterium *Myxococcus xanthus*"

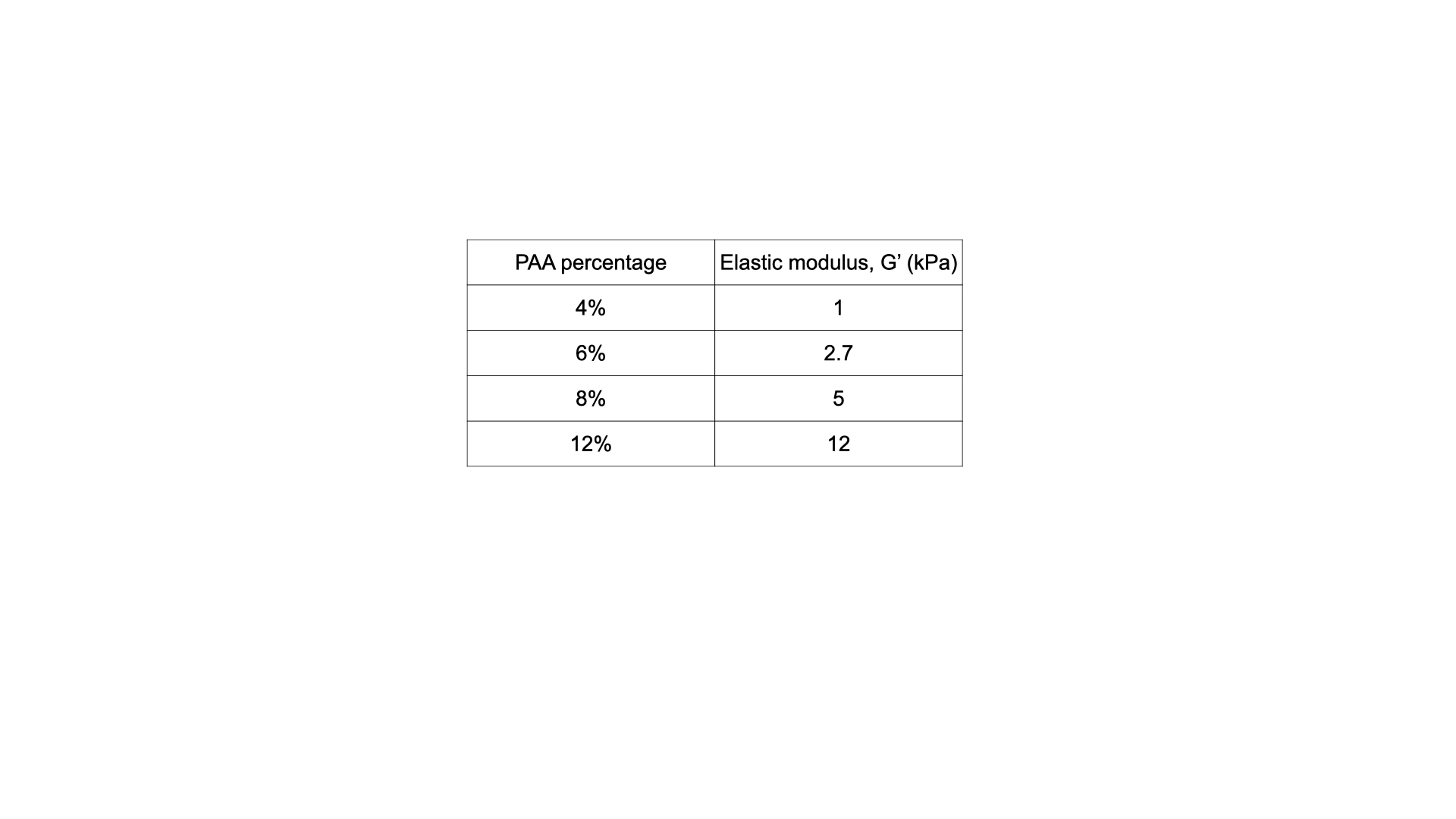

SI. Table.1 Elastic modulus values for varying PAA percentage with 0.15% Bis-acrylamide crosslinker, as measured by oscillatory shear rheology at 5% strain with frequency 1 radians per second.

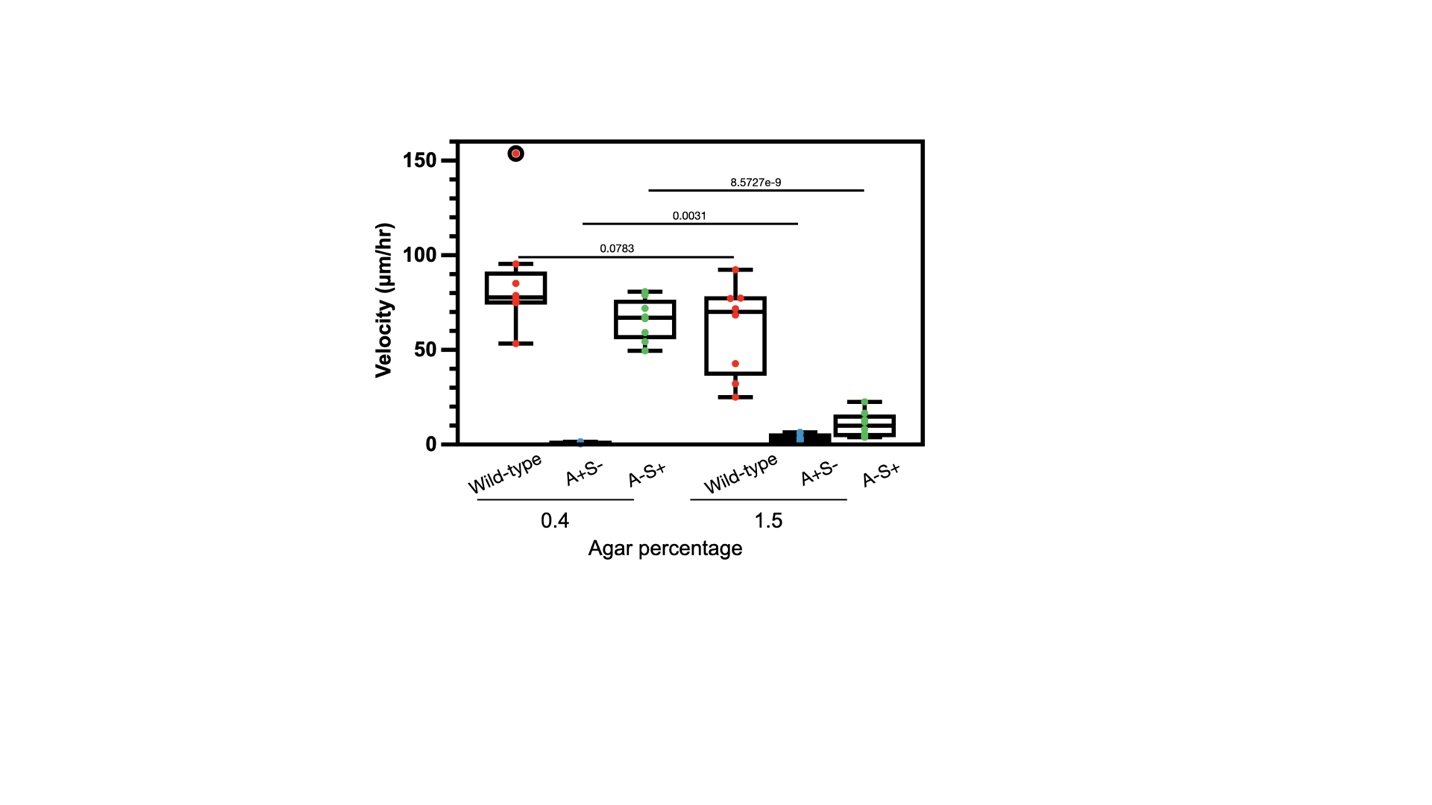

SI. Fig. 1 Velocity of colony expansion at 0.4 and 1.5 agar percentages for *M. xanthus* wildtype (DK1622), A+S-(DK1253), and A-S+(DK1218).
